# High prevalence of Bovine Tuberculosis reported in Cattle and Buffalo of Eastern Nepal

**DOI:** 10.1101/2023.05.21.541660

**Authors:** Nabin Upadhyaya, Nisha Shrestha, Roshan Dahal, Sanjay Kumar Yadav, Rashmi Thakur, Dinesh Aryal, Sujeeta Pokharel, Bibhu Luitel, Rajesh Rajbhandari, Ana M Balseiro, Jose de la Fuente, Christian Gortazar, Dibesh Karmacharya

**Author notes:** Equally contributed as first authors.

## Abstract

Livestock farming, specifically of cattle and buffalo, is crucial to Nepal’s economy, with more than 66% of the population involved in agriculture and animal husbandry. Animal tuberculosis (TB) caused by *Mycobacterium bovis* is a chronic disease that affects these animals and results in economic losses due to reduced milk and meat productivity, fertility and mortality. *M. bovis* also infects humans, non-human primates, goats and other mammals, and can afflict both cattle and buffalo.

Our study is a part of routine surveillance of prevalent diseases, including *M. bovis*, in cattle and buffalo. We collected blood samples (n=400, 100 samples from each district) from selected eastern districts of Nepal. We used a Rapid Bovine TB Test Kit to test these samples for presence of *M. bovis*.

Of the 400 samples collected, 74 animals (18.75%) tested positive for *M. bovis*, with the majority of positive samples coming from cattle (n=71, 17.75%) and only three from buffalo (<1%). Among the screened breeds of cattle and buffalo, Holstein Friesian cattle (HF) (n=43, 58%), Jersey-cross cattle (JX) (n=20, 27%), Local buffalo (n=8, 10.8%) and Murra breed buffalo (n=3, 4.1%) were found to carry *M. bovis*. The majority (50%) of infected animals were between 3-6 years old. Morang (n= 24 positive in cattle; n=1 positive in buffalo) and Jhapa (n= 22 positive in cattle; n=2 positive in buffalo) had the highest prevalence of *M. bovis*, while all the positive cases in Sunsari (n=19) and Udaypur (n=6) were in cattle.

The fact that over 18% of the samples tested positive for *M. bovis* is of great concern. It is critical to thoroughly test animal products from these livestock prior to human consumption. To prevent and mitigate *M. bovis-*related infections in Nepal, a more comprehensive screening strategy coupled with more effective animal husbandry practices needs to be adapted.

## BACKGROUND

In Nepal, the majority of the population (>66%) is involved in agriculture and animal husbandry, and mixed livestock farming is a common practice that often includes poultry and buffalo among other animals (1). Livestock farming in Nepal varies based on ecology and geography (2). In the mountain region, farmers typically rear Yak (Chauri) and sheep, while in the hilly areas cows, sheep, goats, and poultry are common. In the Terai region, popular domesticated animals include buffalo, cows, goats, and poultry (3)

A very common chronic and zoonotic disease caused by members of the *Mycobacterium tuberculosis* complex (MTBC) includes infection by *M. tuberculosis, M. bovis, M. caprae, M. africanum, M. microtia*, and *M. canetti* (4). TB due to *M. tuberculosis* is one of the oldest recorded human diseases, infecting over 10 million people worldwide and causing 1.6 million deaths annually (5). While mainly a respiratory disease, TB can also affect digestive, urogenital and central nervous systems (6). *M. bovis* and *M. caprae*, are the main causes of animal TB and they are found worldwide in both domestic animals (4) and wildlife such as deer, elk, American buffalo, lechwe, and wild boars, among others (7).

*M. bovis* is a chronic and devastating disease that affects both cattle and buffalo in Nepal (8). The disease also poses a threat to humans, non-human primates, goats and other mammals (9). In cattle and buffaloes, the disease generally spreads through contaminated air and milk (10), making it a significant public health concern. The risk of human-to-cattle transmission of the disease is particularly concerning in areas where TB is endemic and its incidence is high (11).

Asymptomatic animals infected with *M. bovis* can shed mycobacteria into unpasteurized milk, contaminated meat, and air which are often the main causes of transmission into humans (12). TB is a chronic disease that affects animal production and reduces reproductive performance in cattle and buffalo (13), resulting in significant economic losses due to increased mortality, reduced milk and meat productivity and reduced fertility (14). Cattle and buffalo are reared in all 77 districts of Nepal, producing nearly 2.5 million metric tons of milk annually (15). The growing number of cattle and buffalo farms, driven by higher demand for milk and milk products, increases the risk of transmission of the disease in humans and animals (8).

In humans, infections caused by both *M. tuberculosis* and *M. bovis* have indistinguishable clinical presentations and similar pathology (9). However, the treatment for these two infections is different, due to the natural resistance of *M. bovis* to pyrazinamide (16). In developing countries, the lack of data on the prevalence of *M. bovis* infection in humans highlights the importance of prevention and control measures, such as mandatory milk pasteurization, adequate meat inspection, and frequent disease monitoring and surveillance in both cattle and buffalo populations (17).

Numerous diagnostic methods have been developed for tuberculosis (TB); sensitivity, specificity, affordability, and portability are some of the key deciding characteristics of good diagnostic tools. The culture method which utilizes Acid Fast Bacilli (AFB) staining, remains the gold standard for diagnosing TB. Several other newer molecular techniques that employ the Polymerase Chain Reaction (PCR) assay are also available (18). In low-income countries, sputum smear microscopy remains the most commonly used diagnostic method. However, diagnosing tuberculosis continues to pose clinical and logistical challenges in resource-limited settings. Point-of-care diagnostics, with high sensitivity and specificity, can be the most practical TB diagnostic tests in resource strapped countries such as Nepal (19).

The main objective of this study is to surveil the prevalence of TB caused by *M. bovis* in cattle and buffalo population of eastern Nepal primarily for two important reasons: i) the eastern districts of Nepal are some of the most important dairy production hubs, ii) and due to their close proximity to the Nepal-India border they experience high transboundary animal movements.

## Methods and Materials

### Study area

Our study sites included areas of Jhapa, Morang, Sunsari, and Udaypur districts [Figure 1]. Diverse breeds of buffalo, including the exotic Indian Murra breed, as well as other indigenous and crossbreeds are found in these areas. These breeds constitute over 35% of the national buffalo population (20). According to livestock statistics, Jhapa (cattle= 247,195, buffalo= 59,930), Morang (cattle= 256,302, buffalo= 52,032 buffalo), Sunsari (cattle= 183,239, buffalo= 39,293), and Udaypur (cattle= 130,021, buffalo= 37,597 buffalo) are among the major livestock-holding districts of Nepal (21).

**Fig 1:**
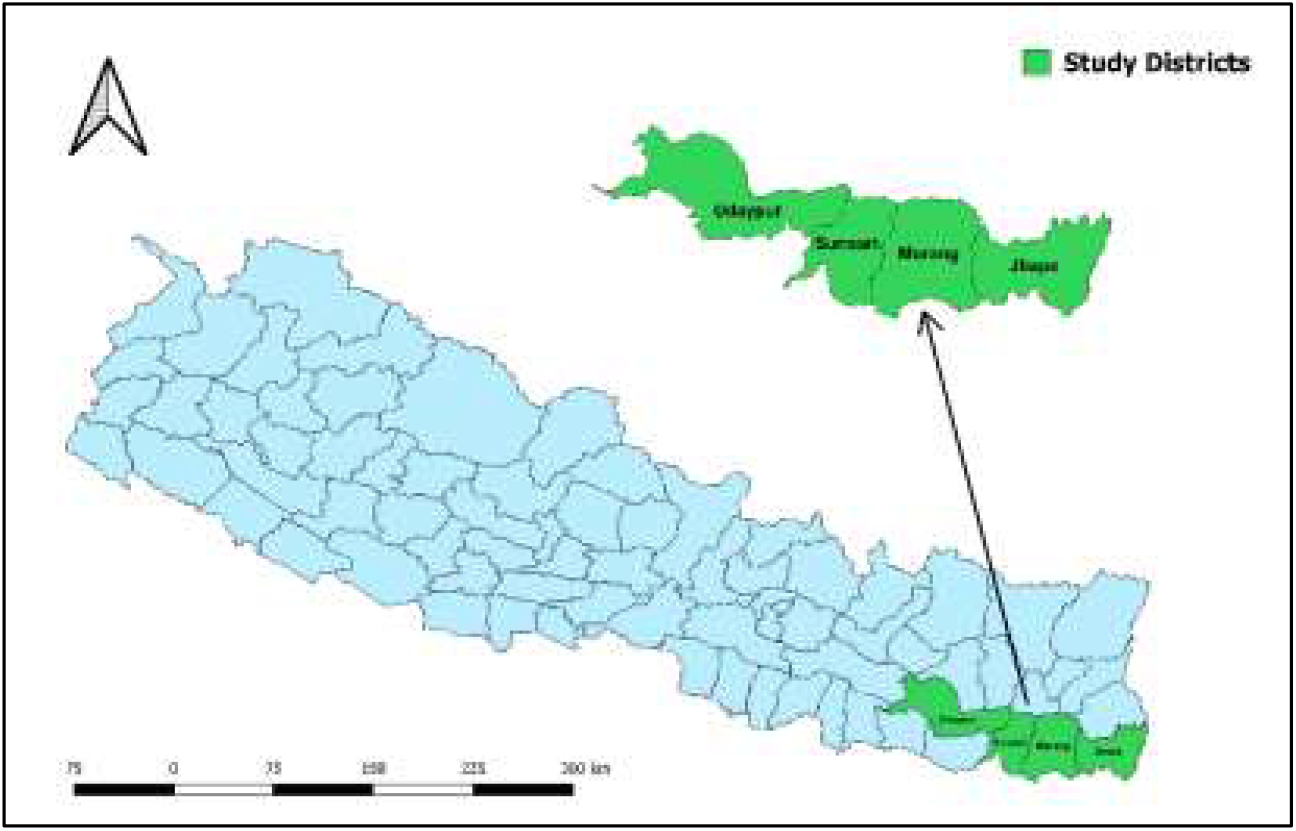
Map of *M. bovis* surveillance study sites in the eastern districts of Nepal.

### Sample collection

A total of 400 animals were included in this study, comprising buffalo (n=9, 2.25%) and cattle (n=391, 97.75%) - these animals were opportunistically sampled at the regional veterinary laboratories. Of the sampled animals, only few were male (n=25, 6.25%) and the rest were female (n=375, 93.75%). Of all the animals tested, majority were Holstein Friesian cattle (HF, n= 225, 56.25%), followed by Jersey-cross cattle (JX, n=115, 28.75%), local buffalo breeds (n=51, 12.75%), and Murra buffalo (n=9, 2.25%).

The Statistical Package for Social Sciences (SPSS Version 20) was used to perform chi-square test on the variables-considering results statistically significant if a two-tailed p-value was less than <0.05.

Blood samples (n=400, 100 samples from each district) were collected from jugular vein using vacutainer clot activator tubes (XLPAC4, Xinle). The serum was separated and was stored in -4 °C freezer. The extracted serum (10ul per test) was then tested for *M. Bovis* using a highly selective recombinant *M. bovis* antigen Rapid Bovine TB test (Bionote; sensitivity 81.7% and specificity 91.4%).

## Results

### Prevalence of Bovine Tuberculosis

Morang (n= 24 positive in cattle; n=1 positive in buffalo) and Jhapa (n= 22 positive in cattle; n=2 positive in buffalo) districts exhibited the highest prevalence of *M. bovis*. In Sunsari (n=19) and Udaypur (n=6) districts, all positive cases were found in cattle (figure 2).

**Fig 2:**
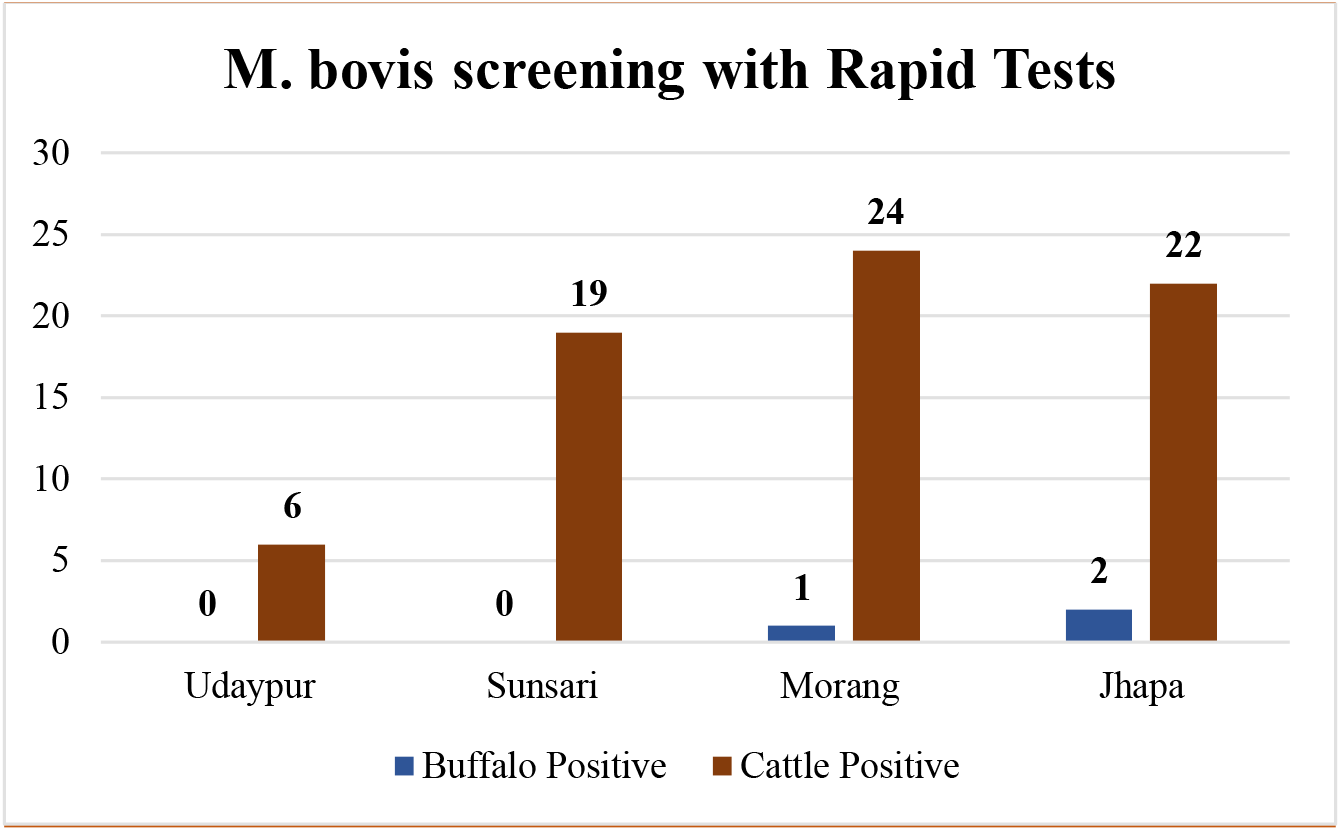
Prevalence of Bovine Tuberculosis in Buffalo and Cattle in the eastern districts of Nepal.

Out of the 400 animals tested, 74 (18.75%) were infected with *M. bovis*, with the majority of the positive samples coming from cattle (n=71, 17.75%) and only three detected in buffalo (<1%). Most of the infected animals were female (n=70, 95%), with only a few males (n=4, 5%) testing positive for *M. bovis* (see Table 1). Among all the screened breeds of cattle and buffalo, Holstein Friesian cattle (HF) (n=43, 58%), Jersey-cross cattle (JX) (n=20, 27%), Local buffalo (n=8, 10.8%) and Murra breed buffalo (n=3, 4.1%) were found to carry *M. bovis* (see Table 2). Half of the infected animals were between 3-6 years old (see Table 3). Our Bi-variate analysis showed no significant relationship between these variables and the trend of *M. bovis* infection.

**Table 1:** Detected *M. bovis* in cattle and buffalo of eastern districts of Nepal.

| Species |  |  |  |  |
| --- | --- | --- | --- | --- |
|  | Buffalo (n= 9, 2.3%) | Cattle (n= 391, 97.8%) | Total | P-value |
| *M. bovis* |  |  |  |  |
| Negative | 6 (1.8%) | 320 (98.2%) | 326 (81.5%) | 0.221 |
| Positive | 3 (4.1%) | 71 (95.9%) | 74 (18.5%) |  |
| Sex |  |  |  |  |
| *M. bovis* | Male | Female | Total | P-value |
| Negative | 21 (6.4%) | 305 (93.6%) | 326 (81.5%) | 0.494 |
| Positive | 4 (5.4%) | 70 (94.6%) | 74 (18.5%) |  |

**Table 2:** Detected *M. bovis* in various breeds of cattle and buffalo.

| **Breed** |  |  |  |  |  |  |
| --- | --- | --- | --- | --- | --- | --- |
|  | Holstein Friesian<br>cattle | Jersey-cross<br>cattle | Local<br>buffalo | Murra<br>breed<br>buffalo | **Total** | **P-value** |
| ***M. bovis*** |  |  |  |  |  |  |
| Negative | 182 (55.8%) | 95 (29.1%) | 43 (13.2%) | 6<br>(1.8%) | 326<br>(81.5%) | 0.630 |
| Positive | 43 (58.1%) | 20 (27.0%) | 8 (10.8%) | 3(4.1%) | 74<br>(18.5%) |  |

**Table 3:** Detected *M. bovis* in cattle and buffalo based on sex.

| **Age** |  |  |  |  |  |
| --- | --- | --- | --- | --- | --- |
|  | 1-3 years | 3-6 years | Above 6<br>year | **Total** | **P-value** |
| ***M. bovis*** |  |  |  |  |  |
| Negative | 163 (50.5%) | 135 (41.8%) | 25<br>(7.7%) | 323<br>(81.5%) | 0.520 |
| Positive | 32 (43.2%) | 36 (48.6%) | 6 (8.1%) | 74 (18.5%) |  |

## Discussion

Our study found that the overall prevalence of *M. bovis* in various species of cattle and buffalo in the selected districts of eastern Nepal was over 18%. This prevalence is similar to another study done in Nepal where *M. bovis* was detected in buffalo (17%) and cattle (16%) using single intradermal cervical test (Jha *et al*., 2007). However, compared to study on *M. bovis* conducted by Pandey *et al*. (22) (prevalence, buffalo= 15.4% and cattle= 13.6%) and Joshi et.al.(23) (prevalence, cattle= 5.78% and buffalo= 9.08%), the prevalence of *M. bovis* in our study was slightly higher. In another study conducted in Nepal, *M. Bovis* was detected in both milk and feces of buffalo and cattle, suggesting increasing contamination of the pathogen in animals and their products in Nepal (24). *M. bovis* cases were detected more in the Jhapa district compared to the Sunsari district, despite Jhapa having a lower cattle population. High transboundary traffic of commercial livestock between Jhapa and India through open border, and lack of animal quarantine facilities might be the possible reasons for this high *M. bovis* prevalence in the district.

There are more cattle than buffalo population in the selected districts, and therefore in our opportunistic sampling we had more cattle samples than buffalo (25). Female cattle are kept for milk production, and hence they outnumber the males-which is reflected in our samples as well (26).

With the high prevalence of *M. bovis* in cattle, there is a significant risk of zoonotic transmission at the cattle-human interface in Nepal. It is important to note that the World Health Organization (2020) has estimated that approximately 117,000 individuals in Nepal are infected with Mycobacterium tuberculosis (27). Given the bidirectional risk of zoonotic transmission, this pose a significant threat to both humans and animals. Since *M. bovis* is the most common causative agent of zoonotic tuberculosis in humans, it is recommended that the relevant authorities implement mandatory testing of cattle using tuberculin skin tests and/or PCR, as well as the pasteurization of milk, as preventive measures.

None of the variables analyzed in our study showed a significant association with *M. bovis* infection. This may be due to the small sample size and/or the selection of non-associated variables. However, a study by Pandey *et al*. (22) showed a significant association between age and *M. bovis* infection in cattle, and we observed a very similar trend-with the *M. bovis* detected more in younger animals. Sex and species were not found to be significantly associated with *M. bovis* infection in our study, which is consistent with the findings of other studies (22,24).

## Limitations of the study

Although the test kit used in this study has high accuracy in detecting antibodies against *M. bovis*, some false results cannot be ruled out. Therefore, additional clinical and/or laboratory tests may be necessary to confirm/validate the results. A relatively small sample size considered in the study, based on opportunistic collection, might not be statistically adequate and hence true picture of actual prevalence of the disease in the animal population might not be very accurate. However, given high prevalence of the pathogen detected in our study, we recommend a more systematic *M. bovis* prevalence study to be carried out in all of the eastern districts of Nepal.

## Conclusion

Based on a rapid test, our study detected *M. bovis* in cattle and buffalo populations in the eastern districts of Nepal- providing a baseline prevalence data. A high prevalence of *M. bovis* of over 18% raises significant concerns for both animal and human health in the region. It is important to thoroughly test animal products, such as milk and meat, from these livestock before human consumption, and a more comprehensive screening strategy along with more effective animal husbandry practices, need to be adapted to prevent and mitigate *M. bovis*-related infections in Nepal.

## Notes

### Competing Interest Statement

The authors have declared no competing interest.

